# The effect of age and sex on the rate of de novo mutations in barn owls

**DOI:** 10.64898/2026.02.17.706346

**Authors:** Alexandros Topaloudis, Anne-Lyse Ducrest, Ana Drago-Rosa, Céline Simon, Bettina Almasi, Alexandre Roulin, Jérôme Goudet

## Abstract

The rate of new mutations is the ultimate source of genetic variation, and thus, a fundamental quantity in evolution. Mutation rates vary across the tree of life, influenced by selection pressures and life-history traits. Within species, mutation rates and types show subtle differences between sexes as well as a positive association with parental age. Most of the direct estimates come from a few mammal species, often primates, and their generality has not been fully realised. Using 33 parent-offspring trios, we investigate the mutation rate in the barn owl (*Tyto alba*) and show that this species has a mutation rate of 5.6 × 10^−9^, matching rates of passerine species with a similar generation time of 2-3 years. Using different phasing methods we show that fathers pass on twice as many mutations as mothers and identify subtle differences in the sex-specific spectra of mutations. We also find that the number of mutations increases with paternal age, bringing direct evidence for germline senescence in birds. While across-species mutation rates and types appear conserved, the potential for within species variation due to age is substantial.

## Introduction

Understanding the rate at which mutations appear can determine both the burden of genetic disease and the pace of adaptation. De novo mutations (DNMs) arise due to mistakes in DNA replication or DNA lesions from mutagenic factors (Baer *et al*., 2007; Seplyarskiy and Sunyaev, 2021), which become incorporated in the DNA sequence due to failure of repair mechanisms (Lindahl and Wood, 1999). These errors are subsequently inherited by daughter cells, leading to the accumulation of mutations with increasing age of a cell lineage, a burden that can, for example, lead to cancer (Moore *et al*., 2021). The mutations that arise in the germline, the subset of cells that will eventually produce gametes, can be inherited by the next generation, introducing heritable variation on which selection can act, leading to evolution.

In the last two decades, accessible sequencing methods have enabled direct measurement of mutation rates through parent-offspring (trio) studies across a broad range of species (Keightley *et al*., 2014; Smeds *et al*., 2016; Pfeifer, 2017; Bergeron *et al*., 2023; Wang and Obbard, 2023; Suárez-Menéndez *et al*., 2023; Zhang *et al*., 2023; Armstrong *et al*., 2025). These studies have revealed that per-generation mutation rates can vary by a factor of 40 across animals. Among the factors influencing this diversity are life-history traits such as generation time and sperm competition, effective population size and genome size (Lynch, 2010; Bergeron *et al*., 2023; Wang and Obbard, 2023). However, so far, estimates of mutation rates for most clades are only represented by a few species, often in one or a few trios of captive individuals. Because the age of an individual and its captivity status can also influence the number of mutations, a more robust approach would involve sampling multiple families of different parental ages from individuals living in the wild. Such study designs can help determine within and between species variation and shed light into the forces driving variation in mutation rates.

Sex and age modulate the rate and type of new mutations introducing variation within species. In mammals and birds, there is a pronounced paternal bias in the mutations inherited by offspring (Kong *et al*., 2012; de Manuel *et al*., 2022; Bergeron *et al*., 2023). In addition, the number of mutations increases with increasing paternal, (Kong *et al*., 2012; Venn *et al*., 2014; Thomas *et al*., 2018; Wang *et al*., 2020, 2022; Versoza *et al*., 2025) and maternal age (Rahbari *et al*., 2016; Goldmann *et al*., 2016). Starting at puberty, spermatogonia undergo continuous mitotic divisions, to produce sperm and replenish themselves. Continuous divisions can lead to accumulating mutations through replication errors although some of these mutations may be instead damage-induced (Gao *et al*., 2016, 2019; Spisak *et al*., 2024). On the contrary, maternal oocytes do not divide post-natally and the increase in DNMs observed occurs exclusively through damage to the DNA. In addition, in humans, mutations from sperm and eggs, have a different nucleotide context (mutational spectrum), due to the different origin of mutations or repair processes which might be sex-specific (Goldmann *et al*., 2016; Jónsson *et al*., 2017; Stringer *et al*., 2020). Most of these findings originate from large human datasets and confirmation of these patterns in other species is lacking.

Birds are a compelling class to study the process of germline mutation as they exhibit substantial variation in life history traits, from short-lived, promiscuous passerines to long-lived, monogamous seabirds. In addition, birds show a lower male bias than mammals (de Manuel *et al*., 2022), but with considerable variation among species (Bergeron *et al*., 2023). Recent work on 18 species (Bergeron *et al*., 2023), found that avian mutation rates differ by 2 orders of magnitude, and that the snowy owl (*Bubo scandiacus*) exhibited the lowest mutation rate among all species analysed (yearly rate of 1.6 × 10^−10^). While there is a strong effect of generation time among species, it remains unclear how the age of the parents influences the rate of germline mutations in birds. In addition, differences in the type of mutations from each sex have not been explored.

We explore these questions using a direct estimate of the germline mutation rate in the Western barn owl (*Tyto alba*) based on 33 parent-offspring trios sequenced at high depth. This large data set, among the largest for any non-human vertebrate, enables us to robustly characterize both the average mutation rate and sex-specific mutation spectra as well as quantify effects of parental age on mutation accumulation. Our findings provide critical insights into the forces shaping germline mutation in a widespread avian predator and contribute to the broader understanding of mutation rate evolution across vertebrates.

## Methods

### Sample selection

A set of 57 individuals (31 males and 26 females) in 33 parent-offspring trios was extracted from a large, monitored population of barn owls from Western Switzerland, for which a field-observation pedigree verified by genomic kinship is available (Topaloudis *et al*., 2025). The choice of individuals was made based on previously available sequences, known age at reproduction and number of offspring of the parents.

### Sequencing & Variant discovery

The DNA of the 57 individuals was obtained from a blood sample taken from the brachial vein during field work, then immediately frozen in liquid nitrogen and stored in a freezer at -80°C until DNA extraction. The DNA was then extracted using the DNeasy Blood & Tissue kit (QIAGEN), quantified using the dsDNA HS Qubit kit or the Quant-it PicoGreen dsDNA Assay kit (Thermo Fisher) and diluted to 6.3 ng/μL with 10 mM Tris-HCl, pH 8.0, in 40 μL for Nextera library preparation and to 1.5 ng/μL in 20 μL for Low-Plex library preparation.

All 57 individuals were previously sequenced (Topaloudis *et al*., 2025). Of these, 52 were re-sequenced to increase the coverage. When possible, we used the same libraries (46 individuals) and generated new libraries from the same DNA extraction for the rest of the samples (6 individuals).

The libraries of 19 samples were previously prepared with PlexWell by Seqwell and sequencing was performed in Illumina NovaSeq 6000. The rest of the libraries (33) were previously prepared with Nextera and sequencing was performed in Illumina HiSeq 4000. For 6 of the 19 Plexwell individuals the old libraries could not be resequenced and new libraries were prepared from a new dilution of the same DNA extraction this time using the Nextera library preparation protocol. All new sequencing data comes from three lanes of a NovaSeq X Illumina sequencing machine. Library preparation and sequencing were handled by the Lausanne Genomics Technology Facility (GTF).

Newly acquired paired-end sequences of 150bp were trimmed to a minimum length of 70bp and adapters were removed using trimmomatic v0.39 (Bolger *et al*., 2014). Reads were aligned to the barn owl reference genome v4 (Machado *et al*., 2022) using BWA-MEM v0.7.17 (Li, 2013). Read groups were added using samtools v.1.19.2 (Danecek *et al*., 2021). Duplicates were marked using GATK v4.5.0.0 (Auwera *et al*., 2013) and *bam* files were merged with previous sequencing data for the same sample. Base quality score recalibration (BQSR) as suggested by GATK ‘best practices’ was performed using a previously published ‘true’ set of SNPs (Cumer *et al*., 2022). GATK’s *HaplotypeCaller* ran for each sample in the barn owl assembly with default parameters and (*-ERC GVCF*). Resulting *gvcf* files were merged per trio using GATK’s *CombineGVCFs* with the parameter (*--convert-to-base-pair-resolution*) which resulted in a file with information about every position in the reference. Variant calling used GATK’s *GenotypeGVCFs* with --*include-non-variant-sites* enabled.

### Filtering

The following filters and steps were applied per trio, in the order presented, to identify putative de novo mutations in the autosomal genome (39 linkage groups):

1. Regions of the genome with poor mappability, annotated repeats or around an insertion/deletion polymorphism (indel) were masked. A mappability mask (Corval *et al*., 2023) removed regions of the genome where less than 95% of the reference fragments (windows of 150bp, step 1bp) did not map uniquely and perfectly. In addition, we masked all sites with an annotated repeat or sites that lie closer than 5bp on either side of the start and end of an indel in each trio.
2. Any variant site with genotype quality (GQ) less than 60 and any non-variant site with a reference genotype quality (RGQ) less than 60 were filtered out.
3. For each sample the mean (*μ̂*) and standard deviation (*σ̂*) of read depth was estimated for variant sites. Any position (variant or not) was removed if it had a read depth less than the larger of 15 or *μ̂* − 2 × *σ̂* . In addition, positions were removed if they had a read depth larger than 2 × *μ̂* or *μ̂* + 2 × *σ̂*, whichever value was smaller. Since all filters above apply to variant and non-variant sites, the remaining sequence is the callable genome (CG) of each trio (see below). The following filters apply only to variant sites and need to be accounted for when estimating the false discovery rate (FDR, β).
4. A modified version of the filters suggested in GATK ‘best practices’ was applied (hereafter technical filters). The threshold values were adjusted to be more stringent, as the original ones are lenient by design. Site were excluded if they failed the following: quality by depth (QD) < 5 or mapping quality (MQ) < 50 or Fisher strand (FS) > 20 or strand odds ratio (SOR) > 2 or mapping quality rank sum test (MQRankSum) < -4 or read position rank sum test (ReadPosRankSum) < -4.
5. Mendelian incompatibilities (MIs) were extracted.
6. MIs were selected where the parents were pure homozygotes (no reads supporting the non-reference allele) and the offspring was a heterozygote. Sites were considered as putative mutations if the offspring had an allelic balance (AB; the ratio of reads supporting the alternative allele over the total number of reads at the site) between 0.3 and 0.7.

Most estimates of mutation rates focus on sites where both parents are homozygous for the reference allele, assumed to be ancestral, and the offspring is a heterozygote. However, we also included sites where both parents were homozygous for the alternative allele and the offspring a heterozygote. We adopted this approach because the reference genome sample came from our study system and the reference allele can not be assumed to be ancestral for all trios.

### Estimating mutation rate

We defined the mutation rate (*μ*) for each trio following (Bergeron *et al*., 2021, 2022; Milhaven *et al*., 2025) as:

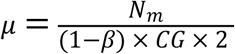

Where *N_m_* is the number of putative de novo mutations after manual inspection, *β* is the false discovery rate (FDR) and *CG* refers to the callable genome, the size of the genome where mutations could be discovered. We estimated each parameter as follows:

- The number of putative de novo mutations *N_m_* was estimated as the number of variants remaining after all filters described above (1-6) were applied. To verify these putative mutations we manually inspected each one in the Integrated Genome Viewer (IGV) v2.19.5 (Robinson *et al*., 2023) following (Keightley *et al*., 2014; Wang *et al*., 2023). For each putative mutation we visualised 2kb around the variant and examined if reads spanning the mutation also contained other alleles not present in the parents in complete linkage disequilibrium. This would suggest a case of missmapping and we labelled such sites as false positives. We also investigated whether parental reads, before GATK’s local realigner, supported the presence of the allele classified as a de novo mutation (DNM). From 536 putative forward (reference to alternative allele) and 44 putative reverse (alternative to reference) after manual inspection we retained 280 forward and 4 reverse mutations as true mutational events. For two true de novo mutations we observed incomplete linkage with an inherited heterozygote nearby, a mosaic pattern, implying a mutation during early development of the offspring. These two sites were considered false positives.
- The callable genome (*CG*), is the number of sites where both parents are homozygotes. These sites were counted after filters one to three were applied (see above). Since all filters up to that step apply to variant and non-variant sites no correction needs to be applied for an FDR. It has been pointed out that the non-variant metric of RGQ is not exactly equivalent to GQ but it is not clear how important the discrepancy is (Pfeifer, 2021). We assumed the effect to be negligible.
- We estimated FDR with an approach adapted from (Bergeron *et al*., 2021, 2023). The approach attempts to account for filters only applied to variant sites because the metrics are not defined for non-variant positions (e.g. allelic balance is only defined in heterozygote genotypes). These filters include the GATK technical filters (which often require 2 alleles) and the AB threshold of 0.3 and 0.7. The underlying hypothesis is that one can correct for these filters by estimating the proportion of true variants filtered out during these steps. For both cases the ‘true variants’ were defined as sites where parents were pure homozygotes for different alleles and the child an inferred heterozygote (depth for each allele > 0). The proportion of these ‘true mutations’ removed after the AB or the technical filters were counted and the sum of these proportions was the estimate of FDR used.

### Sex chromosome

We followed the same approach for the Z chromosome with a key difference. We treated the pseudo-autosomal region (PAR) (Topaloudis *et al*., 2025), and the rest of the chromosome (non-PAR) in two different ways. SNP-calling, filtering and mutation identification in the PAR region was applied exactly as described for autosomal regions. In the non-PAR, females were called as haploids and when applying an individual depth filter in females we halved both the maximum and minimum depth thresholds. For mutation identification in haploid females we did not filter on allelic balance and asked instead that 0 reads supported the reference allele. When estimating the mutation rate in the Z chromosome, we used the following approach to account for the single copy non-PAR in females (multiplied by 3/2 instead of 2):

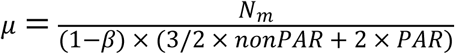

### Phasing de novo mutations

De novo mutations were phased by read-based phasing of the proband using WhatHap v.1.6 (Martin *et al*., 2016). When the DNM was placed in a phase set (i.e. was phased with other nearby heterozygous variants), we looked for phase-informative variants in the phase set to identify paternal or maternal inheritance. Phase informative sites are defined as sites where the proband is heterozygous and at least one of their parents is not.

When possible, (9 individuals), we phased mutations by transmission to a third generation. Phase informative sites (defined as above) were identified in a region of 10kb around the DNM and grand-paternal origin was inferred in the offspring of the proband. Segregation to a third generation was also used to test the Mendelian inheritance of detected DNMs using a binomial test (*binom.test* function in R) of the observed genotypes of the third generation at the mutated site (0 - not inherited, 1 - inherited).

### Mutation spectrum

For each mutation we extracted the two adjacent nucleotides from the reference sequence using bedtools2 (Quinlan, 2014). Mutations were classified according to the nucleotide change that occurred and reverse complements were collapsed (e.g. C > T; G > A). Mutations of the C to T nature occurring in CpG di-nucleotides were separated into their own class. For sex-specific spectra we used the same process but with only the phased mutations coming from each sex. To compare the sex-spectra we used Fisher’s exact test implemented using *fisher.test* in R with simulate.p.value = TRUE and B = 100000 (number of simulations). Multiple testing correction was implemented with p.adjust using the Benjamini and Hochberg method (method=’BH’) (Benjamini and Hochberg, 1995).

### Age effect

Using a generalised linear mixed model with the *glmmTMB* function of glmmTMB package v1.1.13 (Brooks *et al*., 2025), we regressed the phased mutations originating from each sex to the age of the parent of origin (in years), fitting the trio’s callable genome size (in Gb or 10^9^ bp) as a fixed effect and the parent’s id as a random effect. We build a separate model for each sex. The birth year of individuals born in one of the monitored nestboxes is known, and so is their age at reproduction (8 / 16 fathers, 7 / 17 mothers). For the rest of the samples, we can estimate a minimum age by assuming the individual was born at least one year before its first reproductive attempt because barn owls start reproducing as yearlings. Using a generalised linear mixed model, we regressed the inferred phased mutations from each parent to their estimated age. Male mutations were modelled using a Poisson distribution with an identity link. Female mutations were modeled using a zero-inflated Poisson with an identity link, to account for a large number of families with no phased mutations of female origin. Including zero-inflation or not returned almost identical results. Model fitting was tested with DHARMa v0.4.7 (Hartig, 2024). For individuals born in the system the hatch date is known so age can be estimated accurately. For individuals born outside of the studied system, a minimum age can be calculated as one plus the number of years since the first time the individual was observed reproducing, as barn owls can not reproduce before their first winter. This can lead to an underestimation of the age of an individual, thus we ran models using the true and estimated age separately.

Generation time was estimated as the average age of reproducing individuals in the system (Supplementary Figure 1). All statistical analyses and figures were generated using R v4.5.1 (R Core Team, 2025) with the help of the packages, *tidyverse* v2.0.0 (Wickham *et al*., 2019), *ggpubr* v0.6.1 (Kassambara, 2025) and *viridis* v0.6.5 (Garnier *et al*., 2024).

## Results

### Mutation rates

Using 57 individuals in 33 trios, sequenced at an average sequencing depth of 54x (range: 25 - 93x; Figure 1) and stringent variant filtering, we identified 280 forward (reference to alternative) and 4 reverse (alternative to reference) mutations as putative mutational events. On average an offspring carried 8.6 de novo mutations (DNMs) (range 2 - 17, Figure 1). The callable genome in each trio was on average 808 Mb (range 324 - 899) and was influenced by sequencing depth (Supplementary Figure 2). After applying a false discovery rate correction, the average mutation rate per trio was 5.689 × 10^−9^ (range 2.393 × 10^−9^ - 1.042 × 10^−8^). The average age at reproduction in our dataset was 2.727 years bringing the yearly mutation rate to 2.086 × 10^−9^. In 9 individuals (outlined in Figure 1) where segregation of 62 candidate DNMs to the next generation could be tested, we found no evidence of non-Mendelian inheritance of these DNMs using a binomial test.

**Figure 1.**
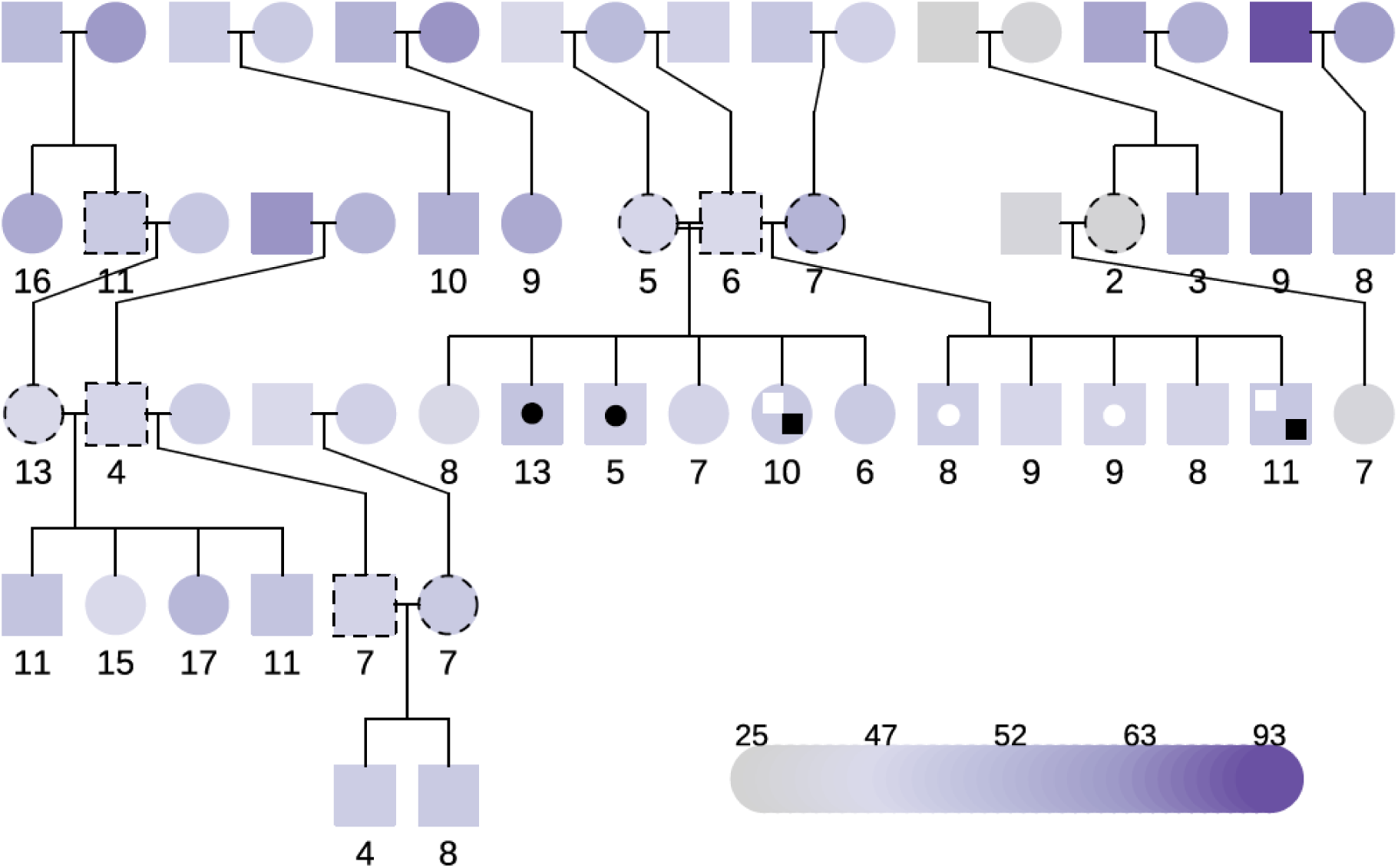
The dataset used in the study. The numbers are the counts of observed mutations after manual inspection. The color shows the average sequencing depth. The dashed outline individuals are those who have both parents and offspring in the dataset and the inheritance of mutations could be tested. The black and white symbols are mutations of maternal (circles) and paternal (squares) origin which are shared between siblings.

Mutations appeared in all autosomal linkage groups (LGs) except one (LG38, size = 7.6Mb). The number of de novo mutations identified per LG correlated strongly with the number of callable sites per LG (Pearson’s correlation, r = 0.7) which in turn scaled almost perfectly (r = 0.99) with a LGs physical length (Supplementary Figure 3). There was no difference between micro- and macro-chromosomes in the number of mutations (β = 0.633, se = 1.5, t = 0.42, p = 0.677). On the Z chromosome we found 4 mutations in two trios, none of which in the pseudo-autosomal region. The per-trio mutation rate for the Z was 5.697 and 6.443 × 10^−8^ and the mutation rate among all trios tested was 5.115 × 10^−9^ within the range of the autosomal estimate.

Of the 284 DNMs, four were shared between 2 siblings each, but were absent from the somatic tissue of the parents (Figure 1). An explanation is that these mutations originated in the primordial germ cells (PGC) of the parents. All four of the putative PGC mutations were C to T transitions. Through read-based phasing we found two of them to be of paternal and two of them of maternal origin. These mutations made up 5% and 10% of the total phased mutations originating from the focal parent(s) respectively. We retained these mutations in downstream analyses as they represent true mutation events that are passed on to the offspring and contribute to the generational mutation rate.

### Mutation spectra

The mutation profile was skewed in favour of transitions (Ti) over transversions (Tv), with a ratio of Ti / Tv equal to 2.5 (Figure 2). The most common class of mutations in the 280 unique identified DNMs, were C to T (including G to A) transitions which comprised 142 (51%) of the total DNMs observed. Of these, 59 (21% of the total) were in CpG dinucleotides. The second most common class was T to C transitions (20%) followed by transversions from C to A (10%). The spectrum was similar to those from multi-trio studies in birds and the one estimated in humans (Supplementary Figure 4).

**Figure 2.**
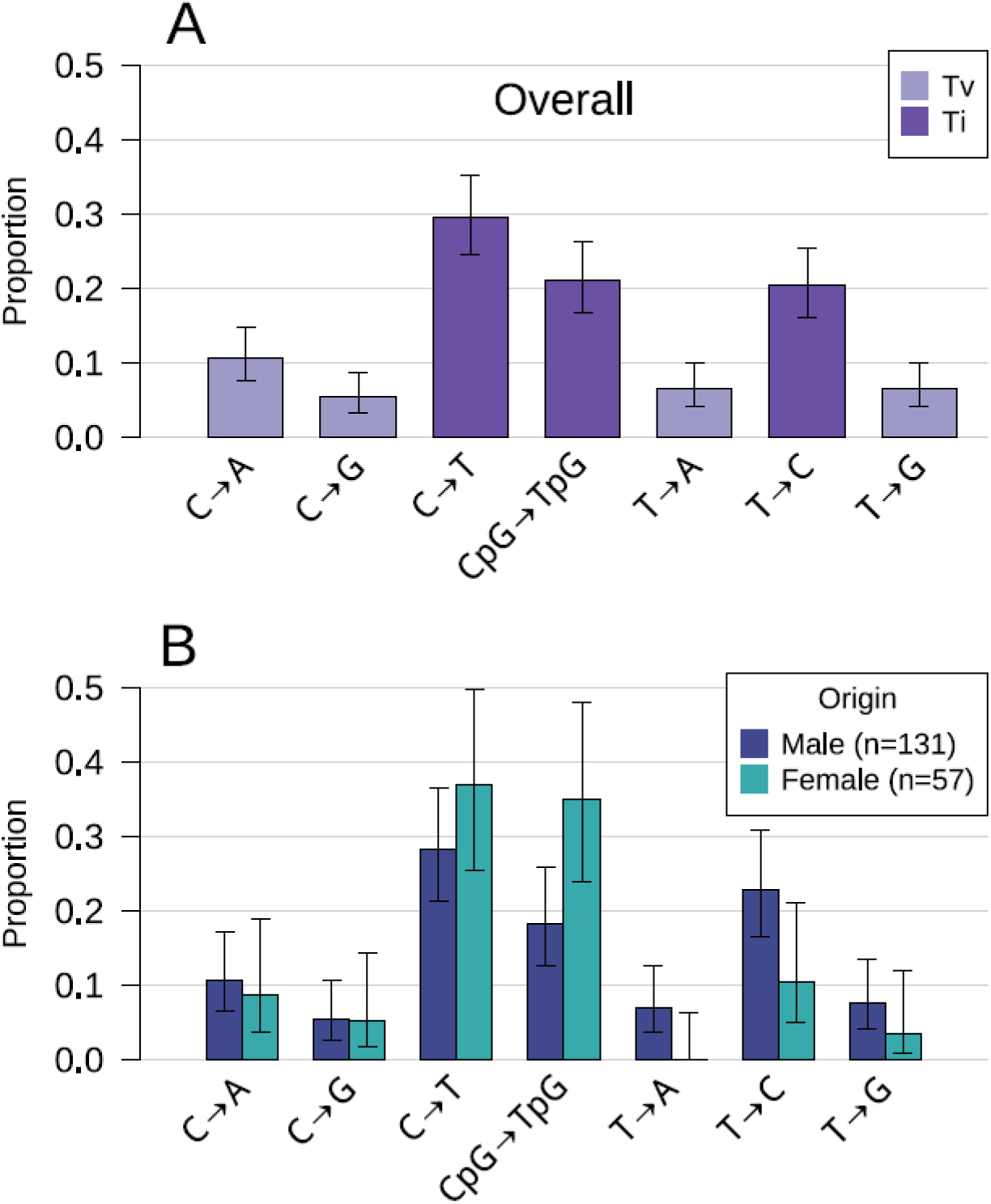
The de novo mutation spectrum. A) Overall mutation spectrum in the 280 unique mutations. B) The proportion of mutations of each class originating in each sex. Ti = transitions, Tv = transversions. Error bars show 95% binomial confidence intervals. Each category includes its reverse complement.

Of the 280 unique de novo mutations we managed to phase 188 (67%). Of these 55 were phased by segregation to a third generation while 175 were phased using read-based phasing and nearby phase informative sites in the focal trio. The average phasing success per trio was 69% (range 17 - 100%). Most of the phased mutations had a paternal origin (131 compared to 57 of maternal origin) bringing the male to female ratio (α) to 2.3. The overall mutation profile was significantly different between the sexes (Fisher’s exact test with simulated p-values: p = 0.026). Maternally inherited mutations tended to be overrepresented in the C to T mutation class, and especially in a CpG context. On the contrary, males passed on a larger number of T to C transitions. As a consequence, the Ti/Tv ratio in females was 4.7 compared to 2.28 in males. No individual class was significantly different after multiple testing correction.

### Effect of age

Paternal mutations explained 29% of the variation in offspring with a significant effect of 0.6 new mutations each year (slope = 0.6, sd = 0.22, z = 2.75, p = 0.006; Figure 3A). Maternal age effects explained 17% of the variation, with an increase of 0.195 mutations per year (slope = 0.195, sd = 0.13, z = 1.5, p = 0.13; Figure 3B) but the slope did not differ significantly from 0. Both estimates correspond to an average callable genome of 804 Mb and refer to successfully phased mutations which are a subset of the total. The models using the full age can be found in Supplementary Figure 5.

**Figure 3.**
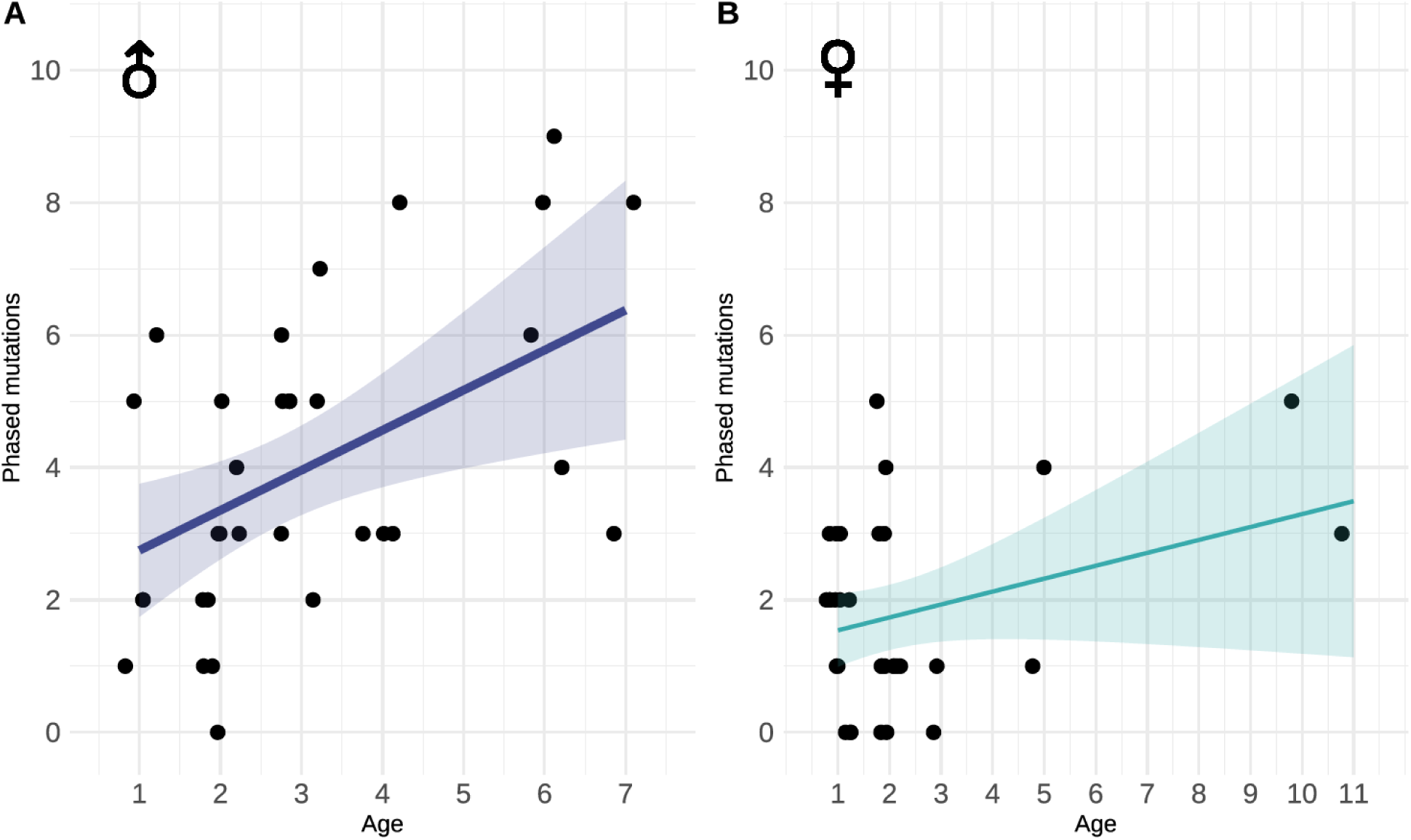
Parental age effect from phased mutations. A) Paternal age effect on the number of mutations inherited from the father. B) Maternal age effect on the number of mutations inherited from the mother. Lines show predicted number of mutations, colored bands show 95% confidence interval.

## Discussion

Using 33 parent-offspring trios we directly estimated the rate of de novo mutations (DNMs) in the barn owl (*Tyto alba*) as 5.69 × 10^−9^. This rate is consistent with other passerine species with similar generation times of ∼2.5 years (Supplementary Figure X) such as the zebra finch (*Taeniopygia castanotis*; g = 2–3 years, μ = 5 × 10^−9^; Prentout et al., 2025; μ = 6.3 × 10^−9^ (Liang *et al*., 2024)), great reed warbler (*Acrocephalus arundinaceus*, g ∼ 2.4 years, μ = 7.6 × 10^−9^; Zhang, Lundberg, et al., 2023), and collared flycatcher (*Ficedula albicollis*, g ∼ 2 years, μ = 4.6 × 10^−9^; Smeds et al., 2016). While direct comparisons among studies may be complicated by methodological differences (Yoder and Tiley, 2021; Bergeron *et al*., 2022), these results reinforce the substantial dependence of mutation rates on generation times and highlight the comparatively low estimated rate of mutation (9.79 × 10^−10^) in the snowy owl (Bergeron *et al*., 2023). This discrepancy could be explained by underlying differences between the two species such as: metabolic rates or whether they choose to reproduce in a year. Alternatively, other extrinsic factors such as the use of a single trio of snowy owls, the captive status of the individuals, the completeness of the reference genome or methodological differences in the pipelines used may also play a role. Using a larger snowy-owl dataset may help to determine if these differences are due to individual level variation or an artefact.

The mutational spectrum we identified can be generated by common mutational processes which act across taxa. The most common type of DNMs we found were changes of CpG to TpG which can occur spontaneously through de-amination when the cytosine is methylated (Cooper and Krawczak, 1989). Because most cytosines in a CpG context are methylated outside of CpG islands, the dinucleotide shows an increased rate towards TpG. The effect is more pronounced when the DNA is found in single-strand conformation, for example in replication and transcription bubbles and in double-strand breaks (Lujan *et al*., 2016; Williams *et al*., 2023; Hinch *et al*., 2023). Further, this mutational signature seems to accumulate with age (Spisak *et al*., 2024) and similar signatures have been found across the tree of life (Campbell *et al*., 2026). In humans (Rahbari *et al*., 2016) and finches (Prentout *et al*., 2025) two mutational signatures make up most of the observed spectrum in the germline. These include the aforementioned de-amination of CpG > TpG and a wide-ranging signature with the hallmark increase in T > C mutations (2nd largest class in our dataset). However, decomposition of mutation spectra into mutational signatures requires either a large dataset of observed DNMs or the use of very rare variants in extensive sequenced datasets, assumed to be very recent and thus influenced by mutational pressure only (Schaibley *et al*., 2013). With the more limited spectrum presented here (7 classes) we observed few differences among species, suggesting a conservation of underlying processes (Supplementary Figure 6). On the one hand this is not surprising given that the forces of endogenous (replication fidelity) and exogenous (DNA lesions) mutagenesis along with repair mechanisms are conserved among species. On the contrary, there is evidence of fine-scale differences in mutation spectra of human populations (Harris, 2015; Mathieson and Reich, 2017; Narasimhan *et al*., 2017) and it remains unclear how much of the observed variation among species, or absence thereof, is due to methodological differences, stochasticity or a true signal.

The two sexes differ in the number of mutations they pass on to the next generation. We found that males pass on twice as many mutations as females, a male bias of 2.3, one of the lowest estimates for birds. Male bias is present in most species where spermatogonia divide continuously (de Manuel *et al*., 2022) as errors accumulate with the rate of mitotic division. In accordance, the male bias is notably lower or absent in seasonally breeding fish where sperm cell production is limited to the mating season (Bergeron *et al*., 2023; Zhang *et al*., 2023). Nevertheless, there is no consensus if male bias is a by-product of the proportion of errors that arise, either due to replication, external damage, or the capacity of the male and female germline to repair such errors (Spisak *et al*., 2024). Since both the frequency of errors and repair efficiency are influenced by epigenetic state, transcriptional activity and the process of recombination, the rate (and type) of mutations in each sex will reflect the combined effects of sex-specific differences in all these fundamental processes during germline fate (Halldorsson *et al*., 2019; Agarwal and Przeworski, 2019; Hinch *et al*., 2023; Wyman *et al*., 2025). As an extension, variation in development of the germline (e.g. methylation state) and life history traits (e.g. sperm competition, cell divisions in each life stage), will impact the evolution of male bias across species. Thus, the cause of mutational male bias involves multiple interlinked processes covering developmental, molecular and evolutionary biology.

In addition to the number of mutations they pass on, we also found that males and females differed in the rate of C > T transitions and the rate of CpG > TpG transitions. An increase in C > T transitions in females is also found in humans (Goldmann *et al*., 2016; Jónsson *et al*., 2017), suggesting conserved differences between sexes either in the origin of mutations or the repair mechanisms employed. In contrast, sex differences in CpG > TpG transitions have not been described in another species. It is known that CpG dinucleotides undergo spontaneous de-amination, an occurrence which correlates strongly and positively with methylation status (Seplyarskiy and Sunyaev, 2021). In humans, the proportion of CpG > TpG transitions is higher in younger females compared to younger males (Jónsson *et al*., 2017). If the same is true in barn owls then it could lead to the observed pattern, as the distribution of female ages in our dataset is skewed towards young individuals (Supplementary Figure 7). For example, 27 of the 33 clutches were mothered by females aged one or two years old (15 of the 33 for males). Alternatively, the difference could be due to a higher overall levels of methylation during oogenesis in birds compared to mammals (Woo and Han, 2024). In addition, it could also point to a large effect of mutagenic recombination, though the mutations due to double strand breaks in humans show an increase of CpT > TpG in the male rather than the female germline (Palsson *et al*., 2025). While the mutagenicity of recombination has not been directly demonstrated in birds (but see Rousselle *et al*., 2019), it is not an unreasonable expectation especially for CpG > TpG transitions which are especially vulnerable at a single strand conformation (Lindahl, 1993). Ideally, these patterns should be verified with a larger dataset of phased mutations from an avian species before concrete conclusions are reached.

The effect of age on the number of mutations was only significant in males, highlighting that paternal age is a key factor of mutational input in birds, mirroring what has already been observed in mammals (Kong *et al*., 2012; Rahbari *et al*., 2016; Goldmann *et al*., 2016; Jónsson *et al*., 2017). In contrast while present, the effect of maternal age was not significant, which may be a result of small sample size and few observations of older mothers as was the case in earlier human studies when the number of trios was limited (Kong *et al*., 2012; Rahbari *et al*., 2016). The parental age effect demonstrated here has important practical implications for avian studies using trio-based mutation rate estimates. In datasets where parental ages are unknown or variable, failure to account for age heterogeneity can introduce substantial variance into mutation rate estimates. In addition, a varying age of the parental cohorts might influence the sex bias inferred since families with older fathers than mothers will exhibit stronger male bias, and vice-versa. Similarly, if the mutation spectrum changes with age as in humans (Jónsson *et al*., 2017; Goldmann *et al*., 2018), the mutational footprint in each sex will also depend on the age of individuals sampled. However, the rate of accumulation of new mutations with paternal age represents only a part of the puzzle as it measures the mutation accumulation post sexual maturity (Ségurel *et al*., 2014; Rahbari *et al*., 2016). Instead a complete picture will require disentangling the relative rates of errors in each life stage along with a knowledge of the number of cell divisions in the germline. Lastly, reproductive performance increases with age in many bird species (Forslund and Pärt, 1995). As a consequence, older males will tend to produce more offspring despite passing on more mutations, a load that could have direct consequences on the fitness of their offspring (Schroeder *et al*., 2015) and result in a trade-off between life-history traits (Monaghan and Metcalfe, 2019).

## Conclusion

While differences in avian mutation rates and types might be discovered as datasets accumulate, we find this rate in barn owls to be remarkably consistent with previous estimates. Species with similar generation times experience similar rates of mutations, suggesting invariable developmental parameters, rates of errors and efficiency of repair. Further we find that mutations originating in paternal germline increased with age.

## Acknowledgments

The authors would like to thank the staff of the Lausanne Genomic Technologies Facility, especially Julian Marquis and Melanie Dupasquier for their assistance in planning and executing the library preparation and sequencing of samples used in this study. They also thank Anna Hewett for improving the manuscript.

## Data availability

Raw sequences will be made available on SRA after acceptance, scripts with all analyses performed can be found on https://github.com/topalw/Mutations.

## Author contributions

A.T and J.G. designed the study. A.L.D. and C.S. performed DNA extraction and dilution. A.T. performed genomic analyses. A.D.R. provided the data on bird age at reproduction. A.H. aided in statistical model design. B.A. and A.R. maintain the barn owl study system. A.T. wrote the manuscript with help from all authors.

