## Supplementary Figure for "The effect of age and sex on the rate of de novo mutations in barn owls"

### Supplementary Material

**Supplementary Material** **1**
Supplementary Figures 2

### Supplementary Figures

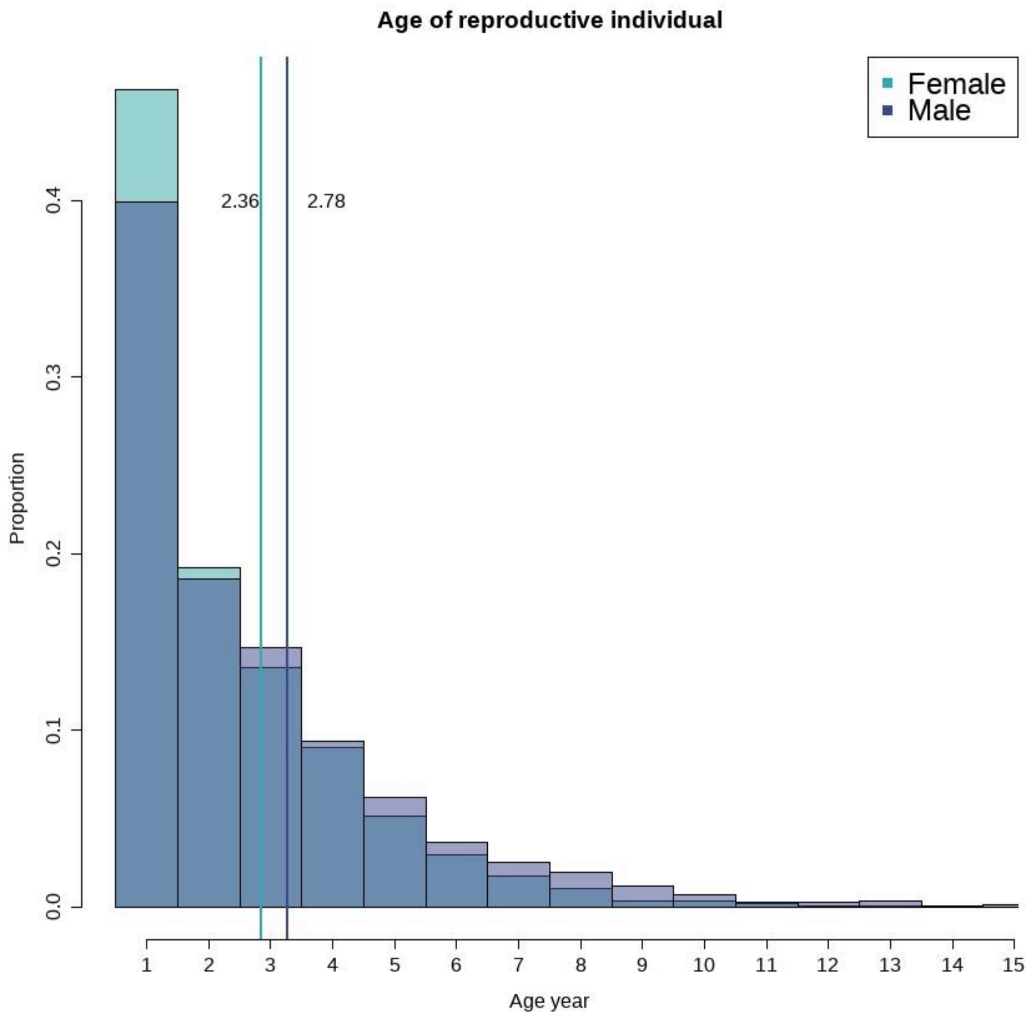


**Supplementary Figure 1. Average age of individuals reproducing in the long term study.** The average age between the sexes is 2.57 years.

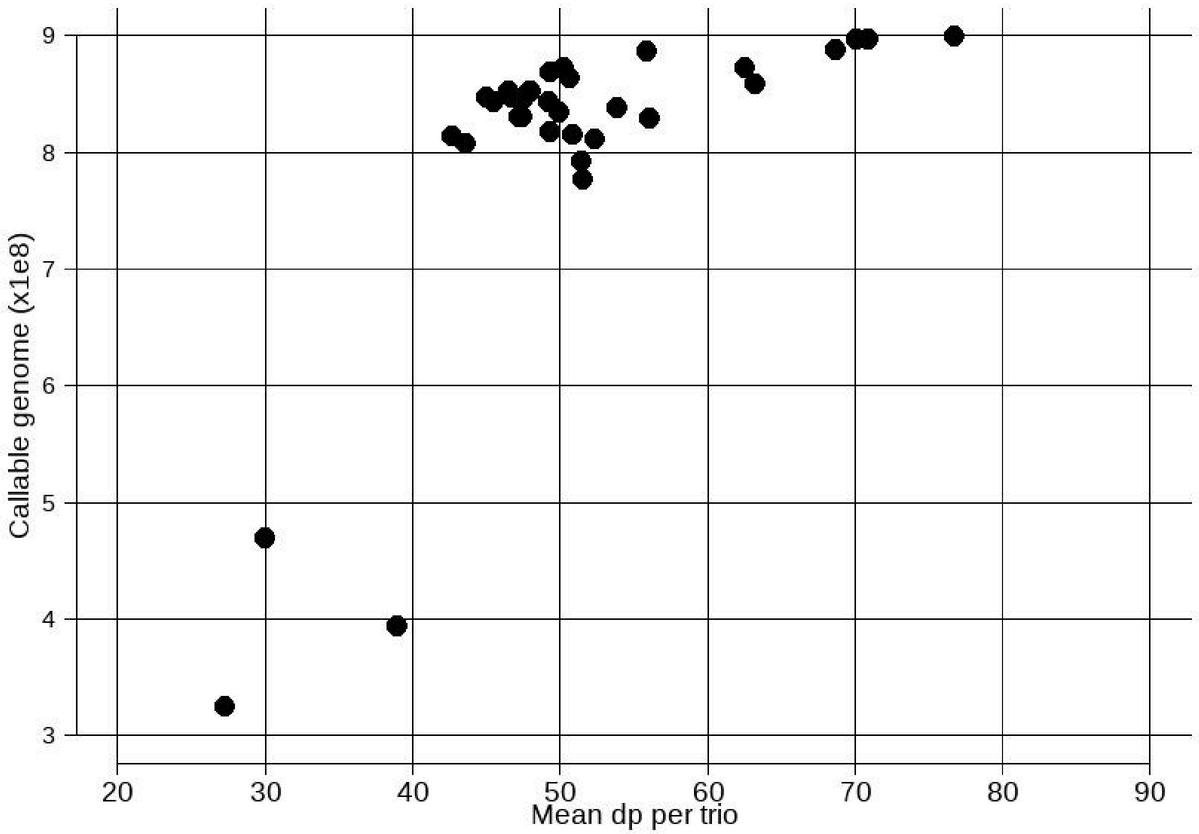


**Supplementary Figure 2. The callable genome(bp) in each trio as a function of average sequencing depth.**

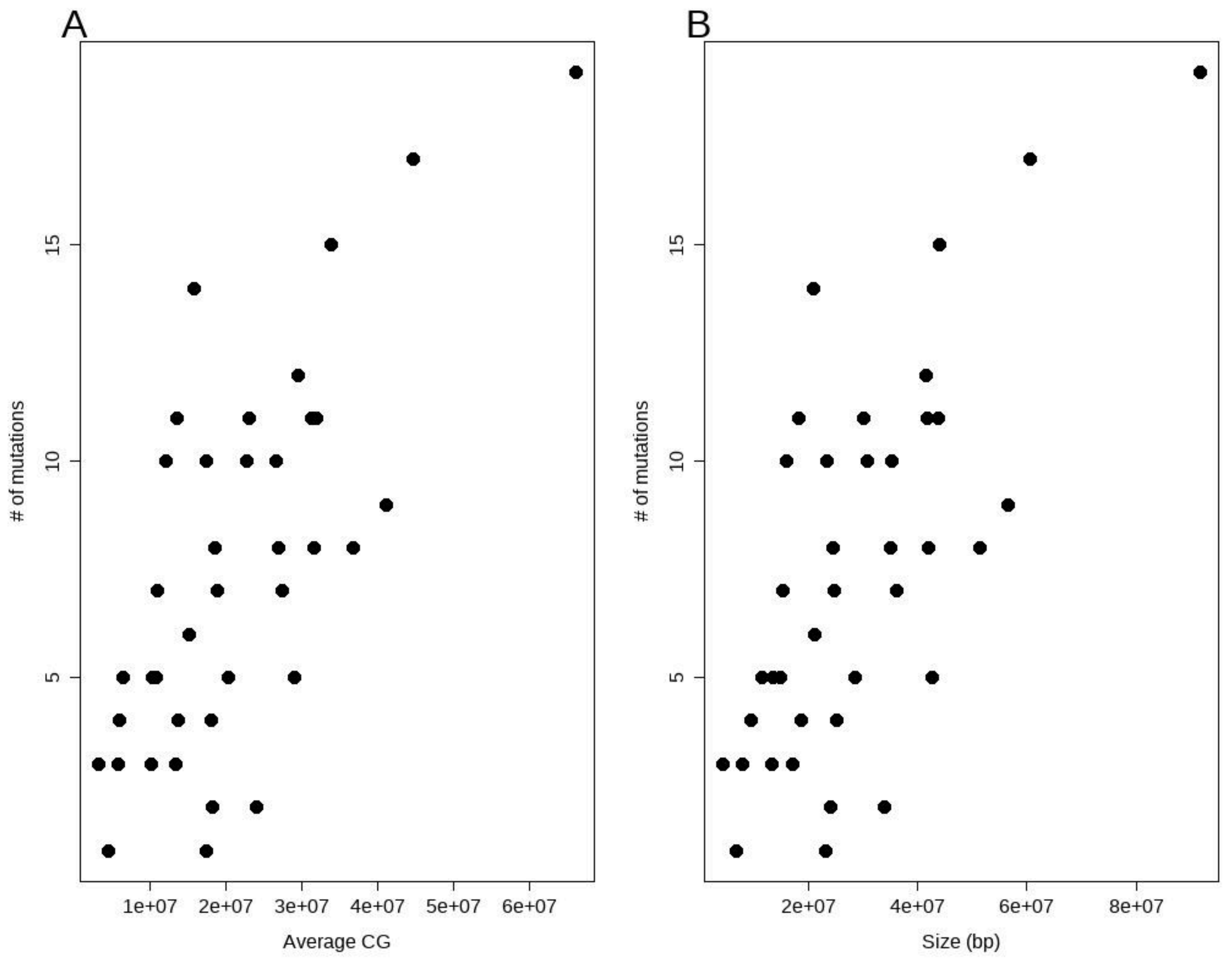


**Supplementary Figure 3. The number of mutations as a function of average callable genome per linkage group (A) or physical length (B).**

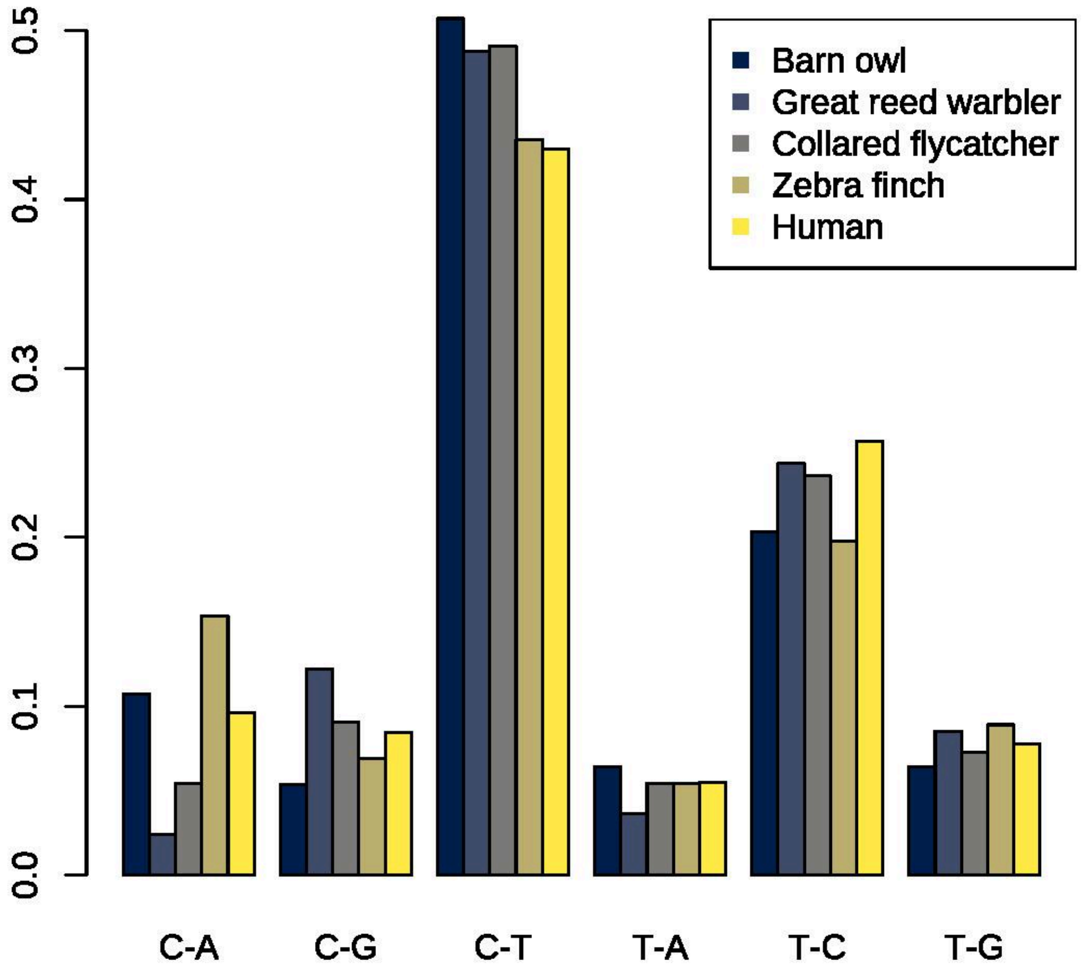


**Supplementary Figure 4. The low resolution mutation spectrum across studies.**
Because some studies did not specify CpG dinucleotides the C>T class contains all C>T mutations. Origin of data: Great reed warbler - (Zhang et al. 2023); Collared flycatcher - (Smeds et al. 2016); Zebra finch (Prentout et al. 2025); Human - (Rahbari et al. 2016)

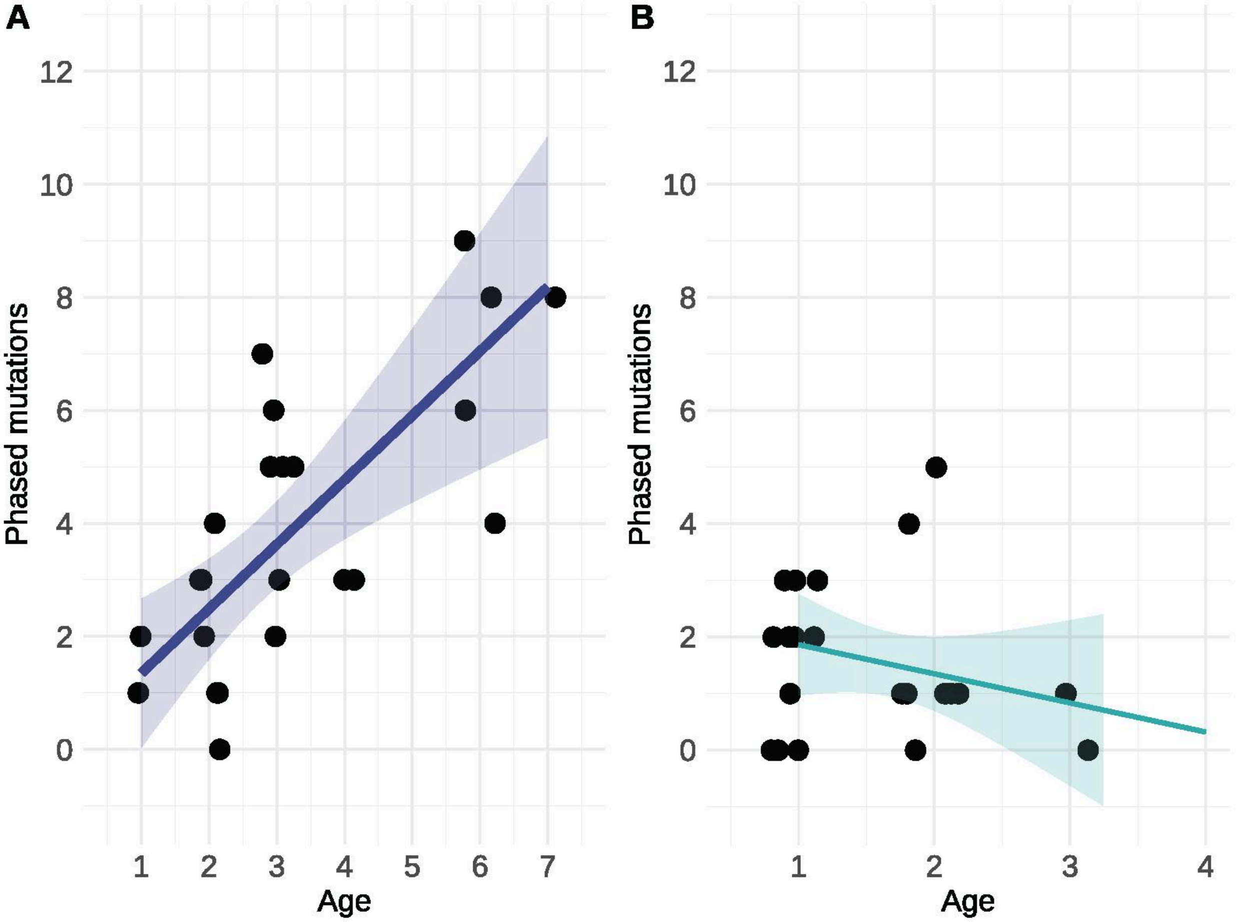


**Supplementary Figure 5. The age effect using only individuals of known age.**
Models as in the main text. Left males (slope = 1.14; z = 3.68; $p < 0.001$); Right females (slope = -0.56; z = -0.95; p = 0.34)

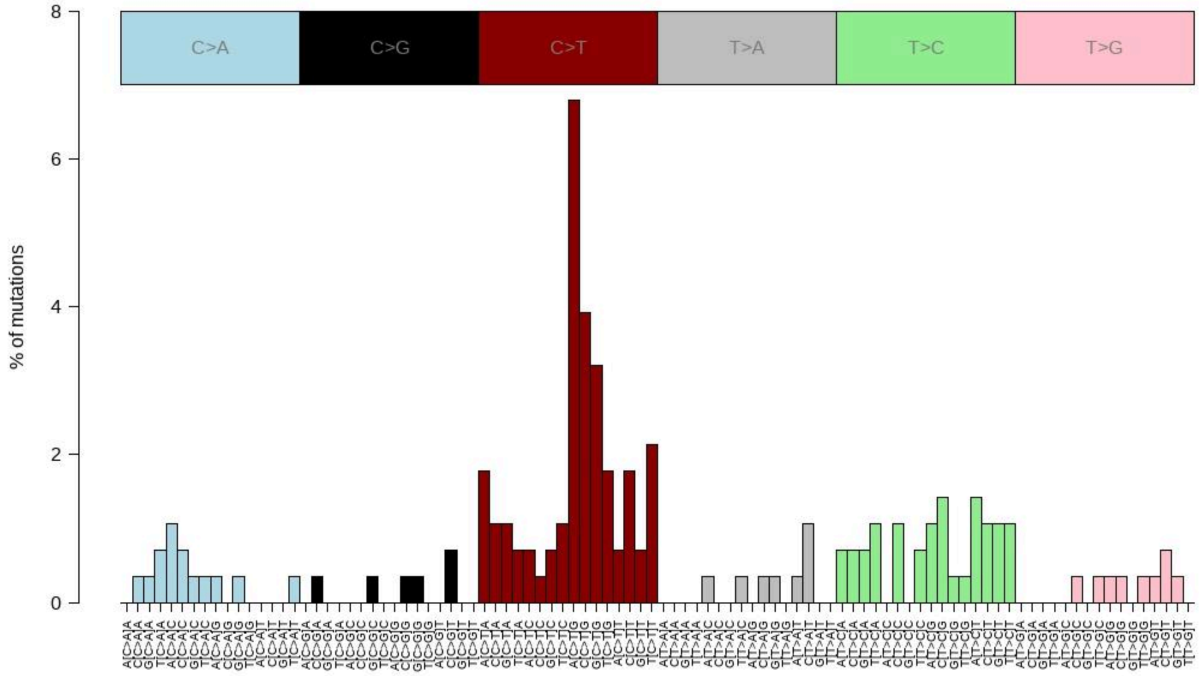


**Supplementary Figure 6. The high resolution mutation spectrum for our dataset.**

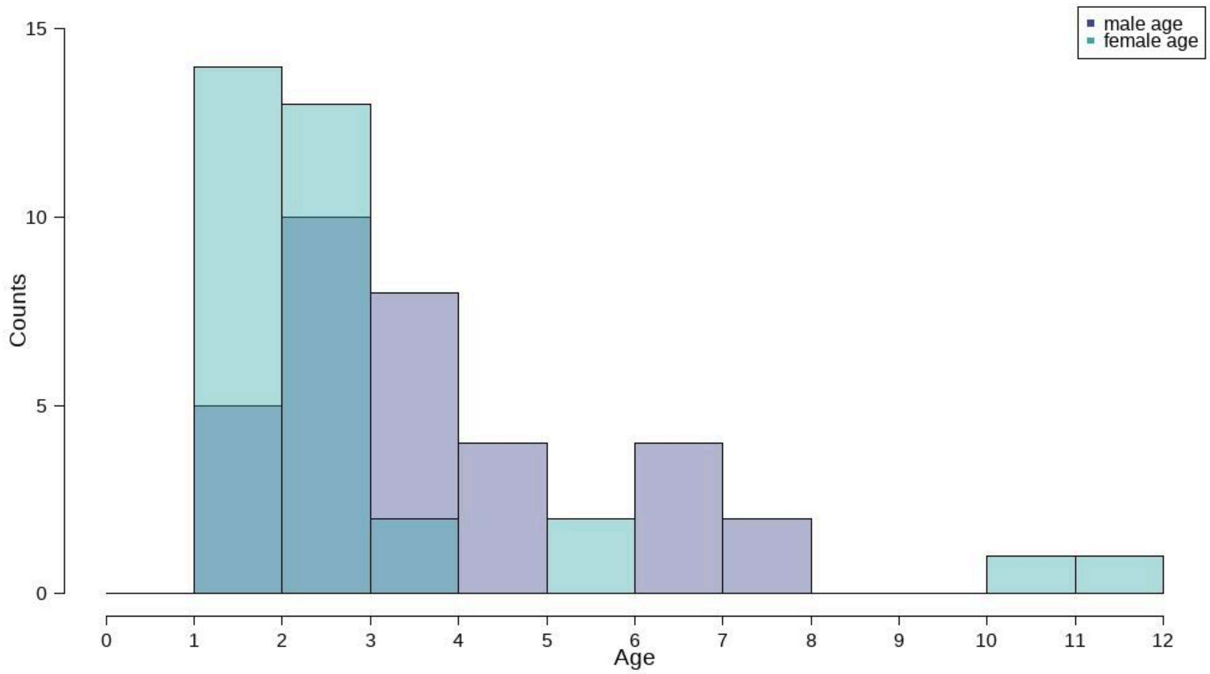


**Supplementary Figure 7. The distribution of age at reproduction in the trios used.**
